# Microinjected dsRNA triggers a robust RNAi response in *Stentor coeruleus*

**DOI:** 10.1101/2025.08.28.672911

**Authors:** Makenna Kuecks, Preeti Arra, Sarah Hoffmann-Weitsman, Makenzie Funk Craig, Mark M. Slabodnick

## Abstract

*Stentor coeruleus* is an emerging model system for the study of morphogenesis and regeneration as well as behaviors like habituation and photosensation. While *Stentor*’s genome has been sequenced, genetic tools are currently limited to RNA interference by feeding. Here we show that microinjection of long double-stranded RNA is an effective method for triggering a robust RNAi response. Additionally, microinjection of *mob1* dsRNA resulted in cell phenotypes appearing 24 hours earlier than by feeding bacterially expressed dsRNA. This work expands the genetic toolkit in *Stentor* when RNAi by feeding is not ideal for an experiment.

## Description

With advances in genome sequencing, research using emerging model systems has increased (Russel et al., 2017). However, even with sequenced genomes additional tool and technique development is a bottleneck when working with organisms that are not widely used and this can severely limit experimental potential in these systems. *Stentor coeruleus* (*Stentor*) is a classical model for unicellular morphogenesis and regeneration that recently had its genome sequenced (Tartar, 1961; Slabodnick et al., 2017). In addition to a sequenced genome, it is also possible to perform RNA interference (RNAi) via feeding cells bacteria expressing long double-stranded RNAs (dsRNA) targeting genes of interest allowing reverse genetic experiments (Slabodnick et al., 2014; Wei et al., 2021). However, one limitation of this method is that it requires completely changing their diet. Cells are commonly fed algae (*Chlamydomonas reinhardtii*), but RNAi by feeding relies on feeding *Stentor* large amounts of either *E. coli* HT115 or *Synechocystis* cyanobacteria for 1-2 weeks in order to trigger the knockdown effect. While this method works well in most situations, for some experiments it is not ideal to drastically alter *Stentor*’s diet and the resulting phenotypes of the knockdown might cause defects in the oral structures and thus the cell’s ability to feed. In *C. elegans* where RNAi by feeding is also possible an effective alternative is the injection of dsRNAs (Fire et al., 1998). Because *Stentor* cells are quite large and easy to inject, we wanted to determine if injection of dsRNA is an effective and viable alternative to RNAi by feeding in *Stentor* as well.

We immobilized *Stentor* in viscous solutions of methylcellulose and then injected them by hand with machine-pulled needles (Fig. 1A). Upon injection, we would observe that cells would swell with the injected solution, forming a clearing in the center of the cell, which we used as a positive marker of injection (Fig. 1A, iii-iv). In very pale cells and in very darkly pigmented cells this clearing was sometimes difficult to observe. Next, we wanted to determine if commonly used storage RNA solutions were toxic upon injection into the *Stentor*. We injected 25 cells with either molecular biology grade water (water), a solution of 0.1 mM EDTA (EDTA), or Tris-EDTA (TE) buffer and observed the cells over a 3-day period counting living cells. We found that there were no significant differences in cell viability between cells injected with water or EDTA on any of the days we observed (Fig. 1B). However, cells injected with TE had significantly lower viability on days 2 and 3 (*p* = 0.045 and 0.018, respectively). In addition to decreased viability, we also noted that many cells were pale in color compared to water or EDTA injected cells. While other buffers should be tested as needed to determine their toxicity, we recommend either water or EDTA be used for *Stentor* injections.

**Figure 1.**
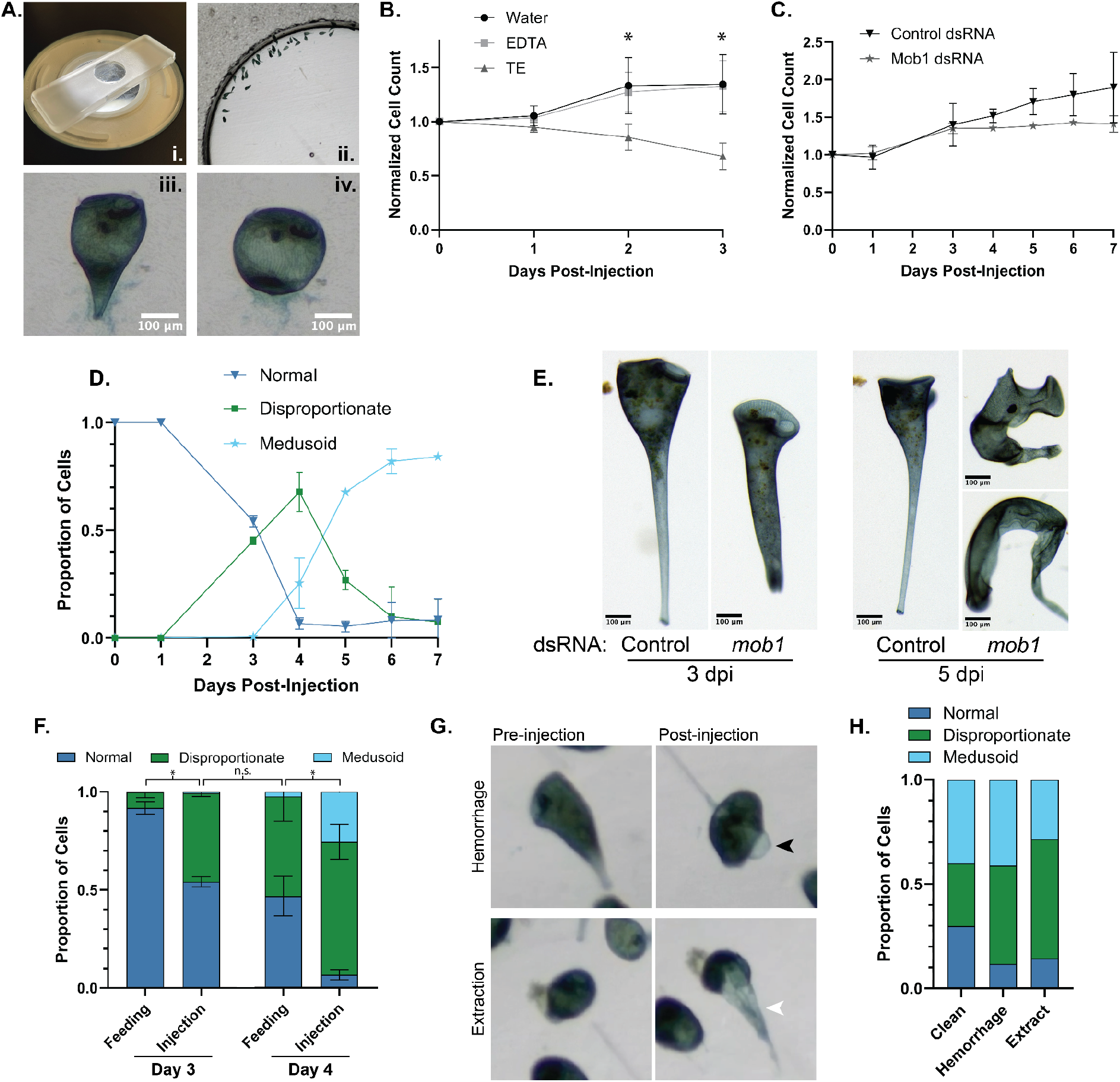
Injection of dsRNA into *Stentor* triggers RNAi. **(A) (i., ii.)** Images showing the injection slide with arranged cells. **(iii**., **iv.)** Brightfield images of a cell before and after injection showing a clearing where the injected material was inserted. **(B)** Cells injected with either water or EDTA show no signs of decreased viability, while cells injected with TE show decreased viability (*) compared to water on day 2 and day 3 (N = 3). **(C)** Cell viability after injecting cells with dsRNA targeting either *lf4* (control) or *mob1*. **(D)** Development of phenotypes in *mob1* dsRNA injected cells over a 7-day period. **(E)** Representative brightfield images of control and *mob1* dsRNA injected cells 3 dpi (left) and 4 dpi (right). **(F)** Comparison of phenotype development after dsRNA injection vs data from previously published RNAi by feeding experiments on days 3 and 5 (Slabodnick et al., 2014; Slabodnick, 2019). **(G)** Representative images showing hemorrhage and extraction after injection and their outcomes regarding viability and phenotype development. **(H)** Cells isolated and binned by injection type showing no significant differences across the samples on 5 dpi (*p* = 0.548).

In order to determine if injected dsRNA is retained and functional inside of *Stentor*, we injected cells with either a control dsRNA encoding the *lf4* gene or dsRNA encoding the *mob1* gene. The *mob1* gene is vital for maintaining proper polarity in *Stentor* and its knockdown results in cells that lose proper proportions along their anterior-posterior axis (disproportionate) and then begin to grow multiple protrusions resembling posterior structures (medusoid). We injected cells with dsRNA encoding either a control gene (*lf4*) or *mob1* on day 0 and followed cells for 7 days post injection (dpi). We didn’t see significant cell death in either sample and the patterns of phenotypes we observed were consistent with previous reports (Fig. 1C-D). Starting on 3 dpi we noted the appearance of many disproportionate cells that lacked the typical trumpet shape while extended (Fig. 1E). These phenotypes progressed into mostly medusoid cells by 5 dpi (Fig. 1E). These data show that with a single injection on Day 0 we can produce the same phenotypes observed with daily feeding of bacterially expressed dsRNA, making experiments less tedious. Additionally, we noted that the appearance of *mob1* knockdown phenotypes seemed to occur rapidly following dsRNA injection.

Therefore, we next compared the timing of the appearance of different phenotypes between dsRNA injection and previously reported data for feeding. While both injection and feeding result in the same patterns of phenotypes, we noted that injection resulted in the appearance of phenotypes about 1 day sooner (Fig. 1F). When comparing dsRNA injected cells to cells fed bacterially expressed dsRNA targeting identical regions of *mob1* the injected cells produced higher number of phenotypic cells by day 3 (*p* = 3.61×10^-20^) and day 4 (*p* = 3.67×10^-27^) while the distribution of phenotypes of day 3 injected cells were similar to day 4 fed cells (*p* = 0.116). This suggests that dsRNA injection would make for more rapid RNAi screening due to the more rapid onset of phenotypes.

Finally, we noticed during injections that cells were not always cleanly injected, with the injection bolus forming a visible clearing inside of the cell’s cytoplasm (Supp. Movie; Fig. 1A, iv). Occasionally, cells would hemorrhage, typically on the side of the cell opposite the injection site, although the bleb that was formed by this hemorrhage appeared to remain in-tact and it was unclear if the injection bolus was leaking out of the cell (Fig 1G, top). Additionally, upon removing the needle from the cell sometimes a large amount of cytoplasm would be extracted from the cell with the needle (Fig. 1G, bottom). We wanted to determine if any of these scenarios resulted in variation in the onset of phenotypes, so we injected cells and binned the cells by the appearance of their injection to follow individual cells and their outcomes. Surprisingly, after 5 days we did not find any significant differences in the onset of RNAi phenotypes in any of the samples (Fig. 1H). This suggests that cell-to-cell variability in the appearance phenotypes is not a consequence of the injection bolus leaking out of the cell due to damage during injection.

Here we show that the use of dsRNA microinjection into *Stentor* is an effective method to trigger RNAi and offers an alternative to bacterial feeding protocols. While additional equipment and skill development are needed, the strengths of this method might outweigh the downsides in many situations. Additional improvements could be developed that would use tracer dyes to follow injected cells but we did not pursue this here. Finally, confirming that injected RNA molecules are retained in *Stentor* and can be processed by the cell suggests that injection of other molecules should be effective. For example, injection of mRNA should result in transient gene expression as it does in other systems and might be easier than delivery and integration of DNA constructs into the nucleus (Ceriotti and Colman, 1995; Hauser et al., 2000). Development of additional tools such as this will continue to strengthen *Stentor coeruleus* as a model system.

## Methods

### Strain and Culture methods

*Stentor coeruleus* cells were originally obtained from a pond on the campus of UNC Chapel Hill (35°54’25.4”N, 79°02’09.3”W) and subsequently kept in continuous culture in the lab. Cells were cultured as previously described with a few modifications (Slabodnick et al., 2014). Briefly, *Stentor* cells were maintained in 50mL glass jars in Ice Mountain spring water (spring water; Primo Brands, Inc.) at room temperature or at 20°C degree in an incubator. Cultures were supplied with 3-4 boiled wheat seeds (Carolina Biological Supply) and fed with *Chlamydomonas reinhardtii* cells (strain CC-124, *Chlamydomonas* Resource Center) twice per week. *Chlamydomonas* cells were cultured as previously described (Harris, 1989) separately in Tris-Acetate Phosphate medium rotating under grow lights and were pelleted and washed in spring water before being fed to *Stentor*.

### dsRNA preparation

Plasmids containing previously validated cloned gene fragments for *lf4* (Control) and *mob1* between opposing promoters for T7 RNA polymerase (Slabodnick et al., 2014) were used as templates for PCR using T7 extension primers (5’-ATAGAATTCTCTAGAAGCTTAATACGACTCACTATAGGG-3’). In vitro transcription was performed using a HiScribe® T7 High Yield RNA Synthesis Kit (New England Biolabs; E2040S) following the manufacturer’s standard protocol. Finally, the dsRNA was resuspended in either water, 0.1mM EDTA, or TE (Tris-HCl pH. 8.0, 0.1mM EDTA) and diluted to 1,000ng/μL for injection. Aliquots were stored at -80°C for up to 6 months.

### Microscopy

General cell handling and injections were performed using a Stemi 508 stereoscope (Zeiss) and images/video were collected using an EXCELIS™ 4K camera (Accu-Scope; AU-800-4K). Images assessing cell shape were recorded using a M205 FCA stereoscope (Leica) equipped with a 1x lens and a DMC5400 camera (Leica).

### Injection setup

Injection needles were pulled on a Pul-1 manual needle puller (World Precision Instruments). Thin wall glass capillary tubes (1.2 OD /0.75 ID) containing a filament were used for injection needles (World Precision Instruments; 1B120F-4). Prior to loading the needle, injection solutions were first centrifuged at 21,000xg for 5 minutes to avoid debris that might clog the needle. Needles were back-filled through capillary action with 1μL of the injection solution that had been Injections were performed using a PLI-100 Pico-Injector (Harvard Apparatus). The pressure was set to 30 psi and the injection time was set to 50 msec. The fused end of the needle was broken off manually using a pair of forceps and the opening was verified by injection into a mineral oil-filled slide visualized under a stereoscope.

### Cell Preparation and dsRNA Injection

For each sample, 30-60 cells were isolated from culture, washed with 750 μL of spring water, and then incubated for 12-24 hours without additional food. Immediately prior to injection, cells were transferred into a solution of 1.5% methylcellulose (Sigma-Aldrich; M0512) dissolved in spring water to slow cell movement and facilitate injection. To maintain order, cells were positioned around the edge of a square-edged circular well for injections (Fig. 1A-ii). All injections were done by hand, holding the capillary holder of the micromanipulator and piercing into the cell membrane with the loaded needle while observing cells under a stereoscope. The injection pedal was then pressed until a visible bubble of injected liquid was visible in the cells (Fig. 1A-iv). The needle was withdrawn and this process was repeated with all of the cells in the well. After injection, 2-3 drops of fresh spring water were added to the top of the methylcellulose to loosen the surface and the cells were gently removed and transferred into 750 μL of spring water. Injected cells were fed immediately and every other day afterward with 20 μL of *Chlamydomonas* and monitored for 7 days.

### Cell viability post-injection

After injection with either water, EDTA, or TE, cells were fed 20μL of washed algae (as described above) and observed once per day for a 3-day period, with the injection day counting as day 0. We quantified viability as the number of living cells in each sample although cells divide during the observation period so this number typically ends above one in healthy samples. Without constant observation, reliably counting cell death is difficult because the shed cortex is sometimes hard to distinguish from cell corpses.

### RNAi Data Collection

Cells were observed every 24 hours after injection to assess viability and cell number, with the exception of 48 hours post-injection for RNAi experiments. Cell viability was scored as alive if cells were able to extend/contract and had active cilia. Cell phenotypes were qualitatively scored as either normal, disproportionate, medusoid as compared to previously reported effects of the *mob1* knockdown (Slabodnick et al., 2014). Representative images of cells were collected on days 3 and 5 post-injection. Cells were carefully positioned and then undisturbed for 1 minute to allow cells to extend before images were collected.

### Data Analysis and Statistics

Data were plotted using Prism (Graphpad Software). For cell viability experiments post-injection, a two-sample T-test was used to compare water against either EDTA or TE conditions. A Chi-Square Test of Independence was used to compare the summed phenotype data between two trials of our injection data and two previously published feeding experiments (Slabodnick et al., 2014; Slabodnick, 2019).

## Acknowledgements

We would like to thank Judy Thorn and Ashley Albright for their comments and suggestions throughout this project and feedback on the manuscript. We would also like to thank the Gerald and Carol Vovis Center for Research and Advanced Study and Health Professions Advising for the work they do to support student undergraduate research at Knox College.

## Funding

We would like to thank the Paul K. and Evalyn Elizabeth Cook Richter Memorial Fund award to Knox College, which supported the purchase of reagents and provided student stipends to complete the work.

## Author Contributions

Makenna Kuecks: Conceptualization, Funding acquisition, Investigation, Methodology, Writing - original draft, Writing - review & editing

Preeti Arra: Conceptualization, Funding acquisition, Investigation, Validation, Writing - review & editing

Sarah Hoffmann-Weitsman: Investigation, Validation, Writing - review & editing

Makenzie Funk Craig: Investigation, Methodology, Writing - review & editing

Mark M. Slabodnick: Conceptualization, Data curation, Formal analysis, Methodology, Project administration, Resources, Supervision, Validation, Visualization, Writing - original draft, Writing- review & editing

## Notes

### Competing Interest Statement

The authors have declared no competing interest.

